# Circulating hemocytes continue to proliferate throughout lifespan in *Daphnia*

**DOI:** 10.1101/2025.10.07.680988

**Authors:** S. C. Cuthrell, L. Y. Yampolsky

**Affiliations:** Department of Biological Sciences, East Tennessee State University, Johnson City TN 37614 USA

**Keywords:** hemocytes, mitosis, Daphnia, ploidy, molting cycle, aging

## Abstract

Daphnia is a classic model organism for the study of circulation and hemocyte biology. Despite the ease of observation and separation of hemocytes, it is not known whether the circulating hemocytes continue to be produced throughout Daphnia lifespan, and if yes, whether they are generated in any specific hemopoietic tissue or represent a reproducing population as known in some arthropods. We detected *de novo* DNA synthesis by means of EdU staining in a significant portion of circulating hemocytes in adult *D. magna*. This portion did not significantly vary among phases of molting/ovary cycle and showed only a slight decrease with age. We did not detect any matching newly synthesized DNA anywhere in any other tissues of adult daphnids, eliminating the hypothesis of the existence of a population of hemopoietic stem cells. Rather, it appears that circulating hemocytes are a reproducing population with mitoses continuing throughout life. Hemocytes showed average ploidy of 4, with a considerable variation around that value indicating that either some endomitoses were also present or a significant portion of hemocytes were sampled in G2. However, while there was no difference in total DNA content measured by Hoechst fluorescence between hemocytes that showed and did not show recent DNA synthesis, among EdU-positive hemocytes there was a positive correlation between Hoechst and EdU fluorescence, indicating that many cells were sampled during S-phase. It is not known whether the total number of hemocytes increases with age or the observed proliferation compensates for losses of hemocytes, as well as whether the EdU-positive hemocytes represent a separate mitotically active stem subpopulation of circulating hemocytes.

## Introduction

*Daphnia* has been one of the first organisms in which hemocytes and phagocytosis have been discovered (Metchnikoff 1884; Tauber 2003) and is currently one of the central model organisms for parasitology research among aquatic invertebrates (Ebert 2005). Yet, we know little about *Daphnia* hemocytes differentiation and dynamics. Although the discovery of hemocytes in invertebrates marked a foundational moment in understanding cellular immunity in all animals, significant differences in hemocyte biology, including hemocyte proliferation, have since been discovered between arthropods and vertebrates. In vertebrates, specifically in mammals, circulating leucocytes, including lymphocytes, are short-lived and non-proliferating, their population replenished by hematopoietic tissues such as bone marrow. Those lymphocytes that are long-lived and capable of proliferating (in response to antigen re-exposure), i.e., memory cells, proliferate largely not in circulation, but in tissues, such as lymph nodes (Abbas et al. 2014). In contrast, in arthropods, besides proliferation in specialized hematopoietic tissues, at least a small proportion of circulating hemocytes show mitotic activity, both background and in response to infection. This has been demonstrated both in insects (Tan et al. 2013; Duressa et al. 2015; Anderl et al. 2016; Boulet et al. 2021) and decapod crustaceans (Sequeira et al. 1996; Gargioni & Barracco 1998; Roulston & Smith 2011; Johansson et al. 2000; Zhao et al. 2022), although Söderhäll and Söderhäll (2022) noted that a significant extent of mitotic activity occurs in the hematopoietic tissues and not in the circulating hemocytes. It should be noted that nearly all data on the dynamics of hemocytes in crustaceans have been obtained in decapods and it is currently not known in what tissues hematopoiesis occurs and whether circulating hemocytes are capable of mitosis in non-decapod crustaceans.

On the other hand, the role of hemocytes in the crustacean immune response, including phagocytosis and melanization, has been well characterized (Johansson et al. 2000; Rowley AF. 2016; Cerenius & Söderhäll 2021). Two types of hemocytes have been described in *Daphnia*: spherical granulocytes and an irregular-shaped amoeboid cells (Auld et al 2010), the later being responsible for cellular immune response. Amoeboid hemocyte circulation significantly increases in response to bacterial (Auld et al. 2010) and *Metschnikowia* yeast infection (Stewart Merrill et al. 2019; Westphal & Stewart Merrill 2022), likely curbing infection by both phagocytosis and defense protein expression. The functions of much more common granulocytes is much less clear, perhaps centered on melanization or on serving as pool of undifferentiated precursors to amoeboid hemocytes. Melanization by the activity of prophenoloxidases expressed by hemocytes plays a role in both parasite encapsulation and wound healing (Terwilliger 2007). In addition, *Daphnia* hemocytes are characterized by a significant expression of cuticular proteins, corroborating the wound healing function (Krishnan et al. 2024; Dua & Yampolsky 2025). It has been shown that, at least in some crustaceans, rates of recruitment circulating hemocytes may increase in sync with molting (Hose et al. 1992), offering both protection against infections during the vulnerable life cycle phase (Söderhäll 2016) and melanization and sclerotization of the carapace (Terwilliger 2007; Alvarez & Chung 2015). Due to the granulocytes constituting a vast majority of circulating hemocytes and because of their possible role in melanization and as precursors of macrophages, the dynamics of this cell type is of special interest and in this study we will be focused on the granulocyte cells, referring to them as hemocytes for simplicity.

This understanding of the dynamics of proliferation and replacement of circulating hemocytes is critical to the understanding of immune response, survival, and longevity in *Daphnia*. A central question would be whether *Daphnia* are post-mitotic with respect to their hemocyte populations, that is, whether or not the circulating hemocytes are terminally differentiated and incapable of cell division.

The current choice of the assay for recent DNA synthesis and thus, for mitotic activity is the EdU (5-ethynyl-2’-deoxyuridine) click chemistry fluorescent staining (Salic & Mitchison 2008). We employed this technique to detect recent proliferation among circulating hemocytes, possible matching mitotic activity in non-circulating stem cells or precursors in potential hematopoietic tissues, and to detect changes in the fraction of hemocytes showing recent DNA replication during the ovary/molt cycle and with age.

## Materials and Methods

*Daphnia* females from the GB-EL75-69 clones (Basel University Daphnia Stock collection, Basel, Switzerland) were sampled from cultures maintained at the standard conditions (20 °C, 12:12 photoperiod, *Scenedesmus acutus* food added in the amount of 1E5 cells/ ml, *Daphnia* density 1 adult per 20 mL of modified ADaM medium, Klüttgen et al. 1994; https://evolution.unibas.ch/ebert/lab/adam.htm). Females were sampled within 24 hours from birth and maintained throughout the lifespan in the same conditions in groups of five in 50 mL plastic inserts placed into 100 mL jars, with jars combined as cohorts depleted due to naturally occurring mortality and to individuals used in experiments, to maintain 5 individuals per 100 mL jar. Neonates were removed every other day and transfer to freshly made medium occurred every 4 days. Five cohorts were started in this manner, 15 – 45 days apart, each replicated 3-fold. Hemocytes proliferation (new DNA synthesis) was assessed by Invitrogen® EdU Click-It kit 3 consecutive batches, every 30 days. The data (logit-transformed fraction of hemocytes showing EdU staining) was analyzed in the entire dataset and, separately, in the longitudinal dataset comprised of the two oldest cohorts sampled in all three batches, and in the common-garden (cross-cohort) dataset comprised of the last of the 3 batches, which included females sampled from all 5 cohorts. Within-cohort replicate was included as a random factor in the complete dataset analysis and both cohorts and replicates nested within cohorts were included as random factors in the longitudinal and common-garden datasets analyses.

EdU exposure was conducted by placing *Daphnia* females individually in 500 uL of ADaM medium containing 100 uM of EdU and 20 uM Hoechst fluorophore for 6 – 24 hours. In a subset of individuals not used in age-specific measurements, the duration of EdU exposure was either 0 (control), or 6, 18, or 24 hours to assess exposure saturation. In all consecutive experiments EdU exposure duration was 6 hours.

To extract hemocytes, at the end of EdU exposure each female (three replicates per replicates, i.e., 9 replicates per cohort, when available) was placed individually into the well of 35 mm, 14 mm well diameter, 1 mm well depth glass bottom dishes (MatTek, Ashland, MA, USA). The EdU solution was removed by pipetting and replaced by 10uL of sterile ADaM medium. Number and developmental stage of embryos in the brood chamber were recorded, with the developmental stage of the brood in the brood chamber to be used to estimate the phase of molting/ovary cycle from ∼24 hours past egg-laying to 94 hours past egg-laying, estimated to be the average timing of the release of the previous brood (following Mittmann et al. 2014 and Toyota et al 2016).

The ventral side of the daphnid was pressed to the wall of the well under a dissecting microscope and an incision was introduced into the dorsal ridge (see Dua and Yampolsky 2025, Fig. 1A) with a tip of a small syringe needle, causing hemocyte-containing hemolymph to bleed. The majority of hemocytes present in the spilling hemolymph are of the spherical “granulocyte” type (Auld et al. 2010). Ten uL of the hemolymph were removed using a 10 uL micropipette with a narrow-bore tip and placed into a fresh glass-bottom dish. No additional permeabilization steps were conducted. 10 uL of 2X staining solution containing Alexa Fluor 488 picolyl azide reagent was prepared according to manufacturer’s protocol to each droplet, followed by 30 min incubation in the dark. At the end of the 30 min period during which the hemocytes remained settled on the bottom of the well, the 20 uL droplet was removed and the cells were washed twice by adding and removing 10 uL of 3% BSA in PBS. The final 10 mL of PBS was retained in the well, and the well was covered with a cover slip, creating a vertical column with 3-4 mm diameter. The hemocytes were then photographed under a Nikon Labophot 2 microscope (10X lens) using UV-1A filter cube (excitation 360-370 nm, dichroic mirror 380 nm, and longpass barrier filter with 420 nm cut-on) for Hoechst staining and with a B-2A filter cube (excitation 450-490 nm, 505 nm dichroic mirror and 520 barrier filter) for EdU / Alexa 488 staining. Hoechst co-staining is necessary to exclude non-specific staining of particles other than nuclei, such as lipid droplets. Nuclei were counted in four non-overlapping view fields in each droplet, and the number of Hoechst- and Alexa-stained nuclei was recorded.

**Fig. 1.**
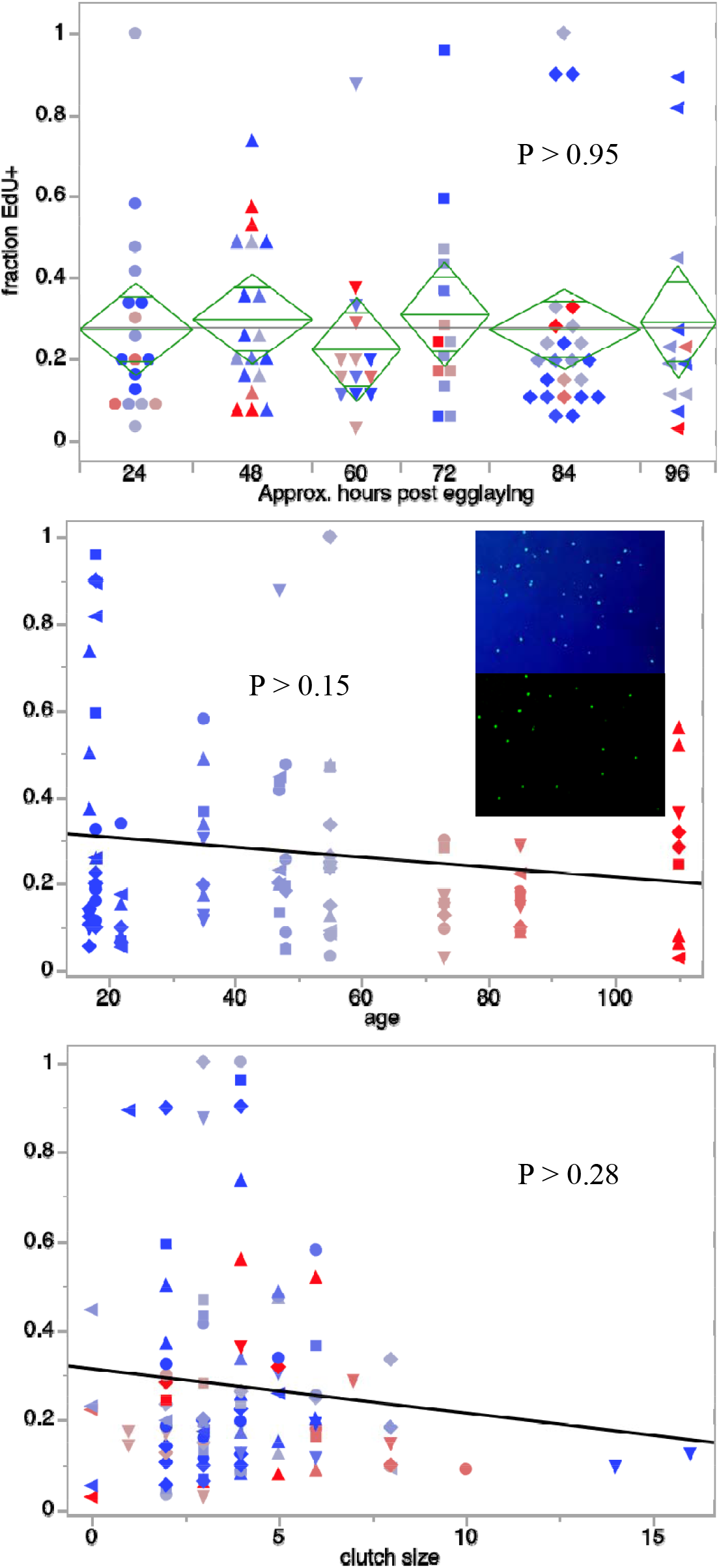
Fraction of hemocytes showing recent DNA synthesis (EdU positive) as a function of ovary cycle phase (hours post egg laying, A), age (days, B), and clutch size (C). ANOVA and linear regression P-values shown. Colors, blue to red, indicate age. Symbols indicate ovary cycle phase (A). See Table 1 for full statistical analysis. Inserts on B: hemocyte nuclei stained with Hoescht (top) and with EdU-specific Click-It Alexa (bottom) fluorescent stains. P-values from a single variable linear regression, see Table 1 and Supplementary Table S1 for complete statistics.

In order to detect EdU staining in other tissues *Daphnia* were treated in the same manner, except that after EdU exposure, adult females were permeabilized with 0.5% Triton X-100, followed by washing twice with 3% BSA in PBS. Juveniles were included in each batch as positive control, as mitotic activity in juvenile *Daphnia* is easily observed in a variety of tissues (Beaton & Hebert 1994).

**Table 1.** Results of REML ANOVA of the effects of age, ovary/molting cycle phase (hours since egg-laying, an ordinal variable), and their interaction on the portion of hemocytes showing EdU staining (combined data). Replicate sub-cohort included into the model as a random variable. See Supplementary Table S1 for the same analysis of longitudinal and common garden datasets separately.

| Source | DF | DFDen | F Ratio | Prob > F |
| --- | --- | --- | --- | --- |
| age | 1 | 62.0 | 0.423 | 0.52 |
| cycle stage | 5 | 51.3 | 0.507 | 0.77 |
| age*cycle stage | 5 | 53.8 | 0.694 | 0.63 |
| clutch size | 1 | 64.5 | 1.151 | 0.29 |
| age*clutch size | 1 | 53.1 | 1.831 | 0.18 |
| cycle stage*clutch size | 5 | 68.1 | 0.559 | 0.73 |
| age*cycle stage*clutch size | 5 | 14.3 | 1.673 | 0.20 |

**Table 2.** The effect of DNA content, measured as background-subtracted total Hoechst fluorescence on EdU fluorescence (also background-subtracted) among hemocytes scored as EdU-positive and EdU-negative. REML ANOVA with images as a random factor. See Fig. 3B for graphic representation.

| Source | DF | DFDen | F Ratio | Prob > F |
| --- | --- | --- | --- | --- |
| DNA content | 1 | 30.0 | 11.731 | 0.0018 |
| EdU status | 1 | 28.1 | 206.159 | <0.0001 |
| EdU status * DNA content | 1 | 29.2 | 10.142 | 0.0034 |

To assess ploidy of hemocytes and other nuclei, *Daphnia* were placed for 2 hours in a 20 uM Hoechst solution, hemocytes extracted as above, and hemocytes, as well as nuclei in antenna-2 muscles, gut, carapace, labrum, and epipodites were photographed as described above. The total amount of DNA in each nucleus was then determined by calculating total background-subtracted fluorescence emitted from the nucleus area. To assess relative amount of Hoechst fluorescence in hemocyte nuclei showing EdU / Alexa 488 staining vs. those that do not, an identical protocol was used, except that hemocytes were photographed under a 20X lens in each fluorescence channel. Hemocytes from four randomly chosen images were analyzed in the following manner. An identical circular region of interest was selected with the central location of the hemocyte nucleus and a histogram of fluorescence intensities was recorded, along with visual identification of EdU-positive nuclei. Then, in each channel, the signal emanating from the nucleus was defined as the area with background-subtracted fluorescence 3 standard deviations of background signal or above over zero. Total fluorescence emanated from these regions in the blue and green emission channels were recorded as, respectively, total DNA and EdU fluorescence.

## Results

### Hemocytes showing recent DNA replication observed in females of all ages

We observed a significant portion of hemocytes positive for EdU staining in females of all ages and reproductive status. The fraction of EdU-positive hemocytes showed a rapid saturation over the duration of EdU exposure from 6 to 24 hours (Supplementary Fig.1). All other experiments were conducted with 6-hours exposure. The fraction of EdU-positive hemocytes did not differ among females in different phases of the ovary/molting cycle, carrying broods of different sizes, or significantly change with age (Fig. 1, Tables 1, S1).

We did not observe any EdU-positive nuclei anywhere in adult females to match the observed numbers of recently proliferated hemocytes (Supplementary Fig.2), with occasionally observed pairs of EdU-positive nuclei indicating that this lack of signal is not caused by problems with staining or permeabilization. This suggests that circulating hemocytes are a mitotically active population which is not replenished from stem cells located elsewhere in *Daphnia* tissues.

### Hemocytes’ ploidy

Consistent with previous data on somatic polyploidy in *Daphnia* (Beaton & Hebert 1989, 1999) different tissues showed a range of ploidy estimates (Fig. 2). Assuming that the lowest total Hoechst fluorescence (from nuclei in antennae muscles) correspond to diploid nuclei, most hemocytes measured appeared to have ploidy of 4 or 8, similar to that of the gut cells and lower than epipodite and labrum highly endopolyploid cells.

**Fig. 2.**
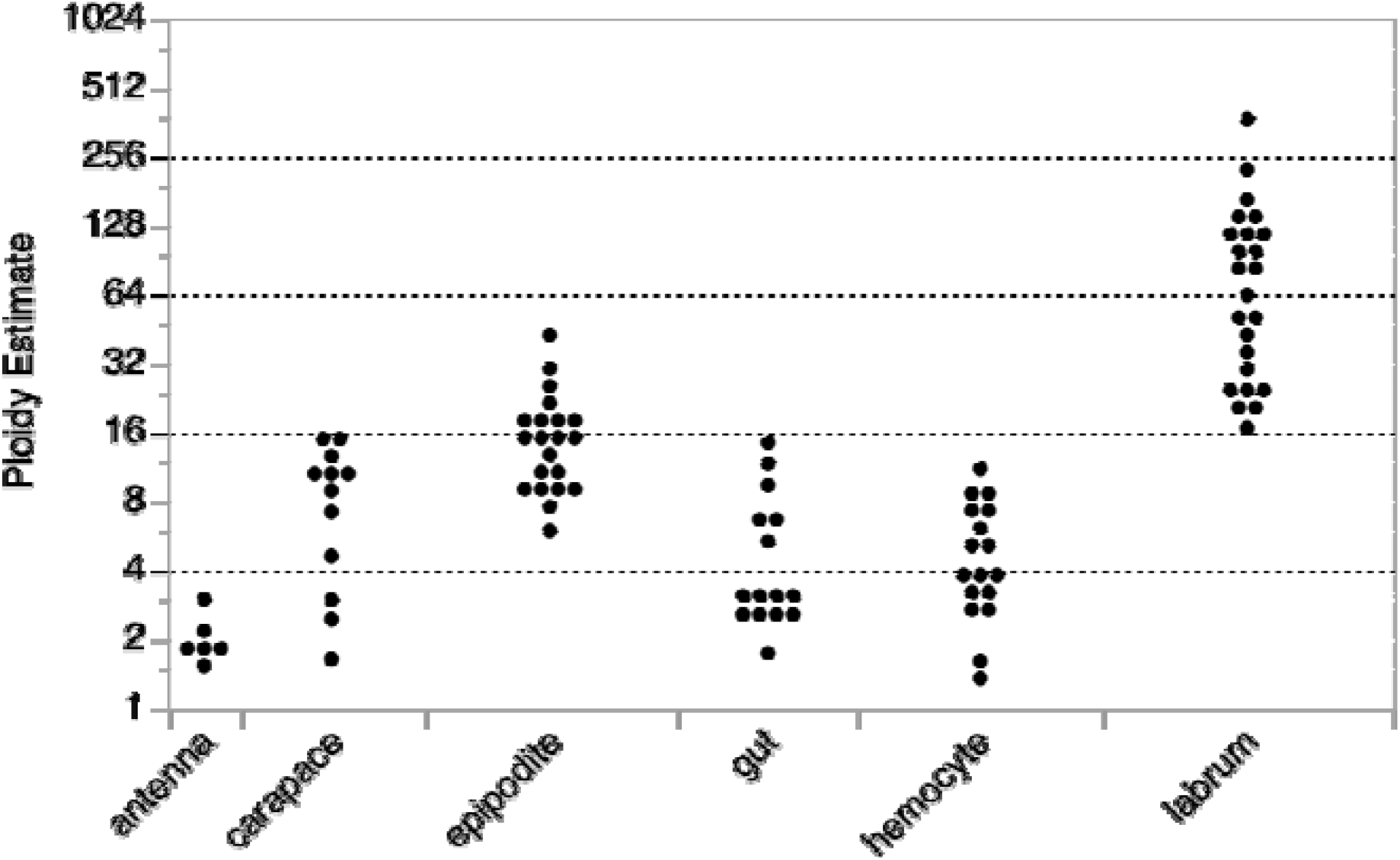
Ploidy estimate in 6 tissue types of *Daphnia magna*.

### Comparison of DNA content in EdU positive and EdU negative hemocytes

Although hemocytes do show variation in ploidy around 4, there were no differences in total DNA content between recently produced and older hemocytes (Fig. 3A). Furthermore, there was a higher variance in EdU-positive than in EdU-negative cells (F_9,23_=3.88; P < 0.0042). Both observations are consistent with the hypothesis that the observed DNA synthesis is associated with cell proliferation and not a manifestation of endomitosis. Furthermore, among EdU-positive nuclei there was a positive correlation between EdU fluorescence intensity and total DNA content (Fig. 3B), indicating that the duration of S-phase was comparable to the 6-hour exposure time. Relative shortage of nuclei with high DNA content, but low EdU fluorescence indicates that the G2 phase is, in contrast, short relative to the duration of exposure. Yet, we cannot completely exclude that some of the EdU-positive hemocytes were endomitotic rather than newly proliferated ones.

**Fig. 3.**
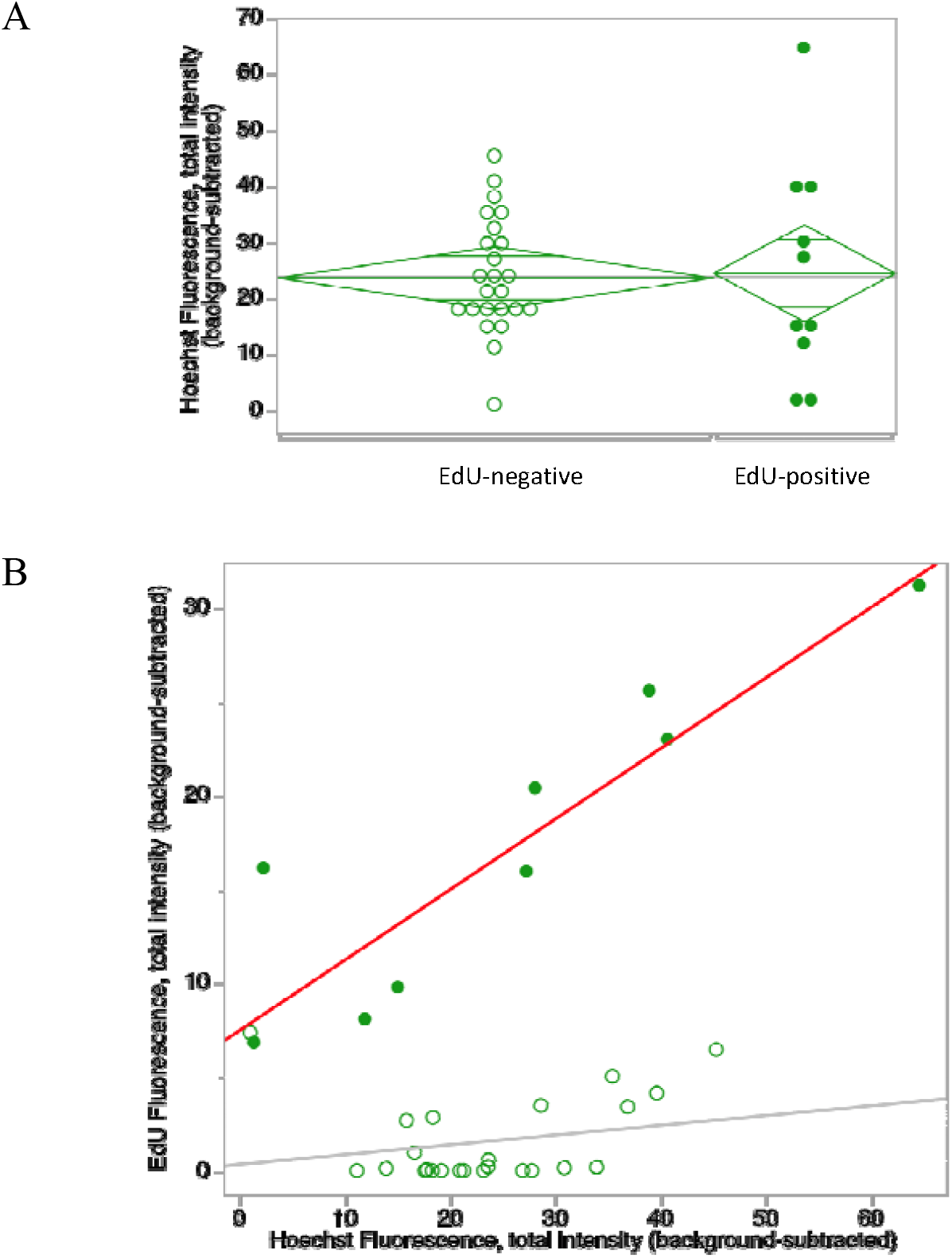
Correlation between DNA content in hemocyte nuclei (measured at total Hoechst fluorescence, background-subtracted) and EdU fluorescence in the same nuclei. A: no difference in DNA content between EdU-positive and EdU-negative nuclei. B: A positive correlation with total DNA content among EdU-positive nuclei.

## Discussion

We observed a consistent occurrence of circulating *Daphnia magna* hemocytes showing recent (within 6 hours) DNA synthesis as indicated by EdU Click-It staining. The fraction of EdU-positive hemocytes did not change along the ovary/molting cycle and did not significantly decrease with age (from ∼30% to about 25% between ages of 20 and 110 days; P>0.15). At the same time, we did not detect any corresponding non-circulating precursor or stem cells with recent DNA synthesis signal, indicating that *Daphnia* circulating hemocytes are a self-replenishing mitotically active population. This is consistent with the observed range of hemocyte ploidy estimates that can be interpreted as the manifestation of some of the EdU-stained nuclei sampled in G1 and some in G2 shortly after mitosis (Fig. 3 B). We can therefore conclude that circulating hemocytes proliferate in *Daphnia* and that this proliferation continues throughout the lifespan. This observation is consistent with similar data in decapods, and is not all too unexpected, despite recent doubts, at least with respect to decapods (Söderhäll & Söderhäll 2022). In fact, the original Metchnikoff’s article describing hemocytes in *Daphnia* contains a drawing of a hemocyte in mitosis (Metchnikoff 1884; Fig. 22).

It is tempting to assume that, given that up to 25% of hemocytes are EdU-positive after 24 hours of exposure and the ovary/molting cycle takes about 4 days at 20 C, hemocytes undergo a complete turnover each cycle. Furthermore, the possible role of hemocytes in sclerotization of cuticulum may imply their increased production at the phases preceding molting (Alvarez & Chung 2015). However, we did not observe any correlation between the fraction of EdU-positive hemocytes and the timing of the ovary/molt cycle. So even if many hemocytes are renewed each cycle, their proliferation appears to be continuous and not linked to molting or egg-laying.

Possible artifactual EdU staining can occur due to either erroneously recording non-specific staining as nuclei staining or due to actual new DNA synthesis detection due to DNA repair rather than replication (Kohlmeier et al. 2013). We did detect up to 5% of green fluorescence particles in some replicates of untreated (no EdU exposure) control (Supplementary Fig. S1), but in other control replicates no EdU-positive particles were detected. This indicates that these replicates may have contained lipid droplets spilled from the *Daphnia* during hemocyte extraction, which were erroneously recorded as stained nuclei. However, in EdU-treated hemocytes, we also cannot exclude that some low-intensity stained hemocyte nuclei represent DNA synthesis during repair rather than replication, so the reported rates of hemocyte proliferation are upper-bound estimates.

Continuous proliferation of hemocytes implies that either the size of circulating population increases throughout the lifespan, or, more likely, that new mitoses compensate for losses of hemocytes. In long-lived decapods, hemocytes’ lifespan can be as long as 2-3 months (Li et al. 2021), and it nevertheless is assumed that recruitment of circulating hemocytes is needed throughout the lifespan regardless of existence of infection to compensate for natural losses (Söderhäll 2016). No data on *Daphnia* hemocyte longevity exists, and, likewise, it is not known whether all circulating hemocytes undergo mitosis periodically or there is a separate stem cell subpopulation that does. We did not investigate the persistence of a population of EdU-positive hemocytes for a prolonged time post-exposure. Such a pulse-chase study might shed light on hemocytes longevity and the possible existence of a circulating subpopulation of mitotically active stem cells (Li et al. 2021).

## Statements and Declarations

### Funding

This study has been funded by Impetus Foundation grant to LYY.

### Competing interests

Authors declare no competing interests.

### Ethics approval, Consent

N/A

### Data, Material and/or Code availability

All data used in the analysis are available in supplementary materials.

### Authors’ contributions

SCC conducted the research and edited the manuscript. LYY designed research, participated in research, analyzed data, and wrote the manuscript.

## Acknowledgements

We are grateful to Marc W Kirschner and Adrian Salic for help with EdU staining application to *Daphnia*, and to A. Catherine Pearson for laboratory assistance. The work was supported by Impetus Foundation grant to LYY.

## Notes

### Competing Interest Statement

The authors have declared no competing interest.

## References

Abbas AK, Lichtman AH, Pillai S. 2014. Cellular and Molecular Immunology, 8th Edition. Saunders, Philadelphia, PA, USA.

Alvarez JV, Chung JS. The Involvement of Hemocyte Prophenoloxidase in the Shell-Hardening Process of the Blue Crab, Callinectes sapidus. PLoS One. 2015 Sep 22;10(9):e0136916. doi: 10.1371/journal.pone.0136916. PMID: 26393802; PMCID: PMC4634603.

Anderl I, Vesala L, Ihalainen TO, Vanha-Aho LM, Andó I, Rämet M, Hultmark D. Transdifferentiation and Proliferation in Two Distinct Hemocyte Lineages in Drosophila melanogaster Larvae after Wasp Infection. PLoS Pathog. 2016 Jul 14;12(7):e1005746. doi: 10.1371/journal.ppat.1005746. PMID: 27414410.

Auld SK, Scholefield JA, Little TJ. Genetic variation in the cellular response of Daphnia magna (Crustacea: Cladocera) to its bacterial parasite. Proc Biol Sci. 2010 Nov 7;277(1698):3291–7. doi: 10.1098/rspb.2010.0772. Epub 2010 Jun 9. PMID: 20534618; PMCID: PMC2981931.

Beaton MJ, Hebert PD. Patterns of DNA synthesis and mitotic activity during the intermoult of Daphnia. J Exp Zool. 1994 Apr 1;268(5):400–9. doi: 10.1002/jez.1402680509. PMID: 8158101.

Beaton, M.J., Hebert, P.D.N. 1989. Miniature genomes and endopolyploidy in cladoceran crustaceans. Genome 32, 1048–1053.

Beaton, M.J., Hebert, P.D.N., 1999. Shifts in postembryonic somatic ploidy levels in Daphnia pulex. Hydrobiologia 394, 29–39.

Boulet M, Renaud Y, Lapraz F, Benmimoun B, Vandel L, Waltzer L. Characterization of the Drosophila Adult Hematopoietic System Reveals a Rare Cell Population With Differentiation and Proliferation Potential. Front Cell Dev Biol. 2021 Oct 13;9:739357. doi: 10.3389/fcell.2021.739357. PMID: 34722521.

Cerenius L, Söderhäll K. Immune properties of invertebrate phenoloxidases. Dev Comp Immunol. 2021 Sep;122:104098. doi: 10.1016/j.dci.2021.104098. Epub 2021 Apr 20. PMID: 33857469.

Dua I, Yampolsky LY. Transcriptional atlas of Daphnia magna. Comp Biochem Physiol Part D Genomics Proteomics. 2025 Mar 31;55:101504. doi: 10.1016/j.cbd.2025.101504. Epub ahead of print. PMID: 40199048.

Duressa TF, Vanlaer R, Huybrechts R. Locust cellular defense against infections: sites of pathogen clearance and hemocyte proliferation. Dev Comp Immunol. 2015 Jan;48(1):244–53. doi: 10.1016/j.dci.2014.09.005. Epub 2014 Oct 3. PMID: 25281274.

Ebert D, 2005. Ecology, Epidemiology, and Evolution of Parasitism in Daphnia. National Library of Medicine (US), National Center for Biotechnology Information, Bethesda MD, USA. Available from: http://www.ncbi.nlm.nih.gov/entrez/query.fcgi?db=Books

Gargioni R, Barracco MA. Hemocytes of the palaemonids Macrobrachium rosenbergii and M. acanthurus, and of the penaeid Penaeus paulensis. J Morphol. 1998 Jun;236(3):209–21. doi: 10.1002/(SICI)1097-4687(199806)236:3<209::AID-JMOR4>3.0.CO;2-Y. PMID: 9606943.

Hose JE, Martin GG, Tiu S, McKrell N. Patterns of Hemocyte Production and Release Throughout the Molt Cycle in the Penaeid Shrimp Sicyonia ingentis. Biol Bull. 1992 Oct;183(2):185–199. doi: 10.2307/1542206. PMID: 29300648.

Johansson MW, Keyser P, Sritunyalucksana K, Söderhall K. 2000. Crustacean haemocytes and haematopoiesis. Aquaculture 191: 45–52.

Kohlmeier F, Maya-Mendoza A, Jackson DA. EdU induces DNA damage response and cell death in mESC in culture. Chromosome Res. 2013 Mar;21(1):87–100. doi: 10.1007/s10577-013-9340-5. PMID: 23463495.

Krishnan I, Yampolsky LY, Petrova K, Peshkin L. 2024. Single Cell Transcriptome Defines Cell Type Repertoire of Adult Daphnia magna. bioRxiv 2024.05.29.596540; doi: 10.1101/2024.05.29.596540

Metchnikoff, E. (1884) Uber eine sprosspilzkrankheit der daphnien; Beitrag zur über den kamp der phagocyten gengen krankheitserreger. Arch. Pathol. Anat. Physiol. Klin. Med. 96, 177–195. 10.1007/BF02361555.

Mittmann B, Ungerer P, Klann M, Stollewerk A, Wolff C. Development and staging of the water flea Daphnia magna (Straus, 1820; Cladocera, Daphniidae) based on morphological landmarks. Evodevo. 2014 Mar 18;5(1):12. doi: 10.1186/2041-9139-5-12. PMID: 24641948; PMCID: PMC4108089.

Roulston C, Smith VJ. Isolation and in vitro characterisation of prohaemocytes from the spider crab, Hyas araneus (L.). Dev Comp Immunol. 2011 May;35(5):537–44. doi: 10.1016/j.dci.2010.12.012. Epub 2010 Dec 22. PMID: 21184777.

Rowley AF. 2016. The Immune System of Crustaceans. Encyclopedia of Immunobiology, 437–453. doi:10.1016/b978-0-12-374279-7.12005-3

Salic A, Mitchison TJ. A chemical method for fast and sensitive detection of DNA synthesis in vivo. Proc Natl Acad Sci U S A. 2008 Feb 19;105(7):2415–20. doi: 10.1073/pnas.0712168105. PMID: 18272492.

Sequeira T, Tavares D, Arala-Chaves M. Evidence for circulating hemocyte proliferation in the shrimp Penaeus japonicus. Dev Comp Immunol. 1996 Mar-Apr;20(2):97–104. doi: 10.1016/0145-305x(96)00001-8. PMID: 8799615.

Söderhäll I, Söderhäll K. Blood cell formation in crustaceans. Fish Shellfish Immunol. 2022 Dec;131:1335–1342. doi: 10.1016/j.fsi.2022.10.008. Epub 2022 Oct 7. PMID: 36216230.

Söderhäll I. Crustacean hematopoiesis. Dev Comp Immunol. 2016 May;58:129–41. doi: 10.1016/j.dci.2015.12.009. Epub 2015 Dec 23. PMID: 26721583.

Stewart Merrill TE, Hall SR, Merrill L, Cáceres CE. Variation in Immune Defense Shapes Disease Outcomes in Laboratory and Wild Daphnia. Integr Comp Biol. 2019 Nov 1;59(5):1203–1219. doi: 10.1093/icb/icz079. PMID: 31141120.

Tan J, Xu M, Zhang K, Wang X, Chen S, Li T, Xiang Z, Cui H. Characterization of hemocytes proliferation in larval silkworm, Bombyx mori. J Insect Physiol. 2013 Jun;59(6):595–603. doi: 10.1016/j.jinsphys.2013.03.008. PMID: 23557681.

Tauber, A. Metchnikoff and the phagocytosis theory. Nat Rev Mol Cell Biol 4, 897–901 (2003). 10.1038/nrm1244

Terwilliger NB. Hemocyanins and the immune response: defense against the dark arts. Integr Comp Biol. 2007 Oct;47(4):662–5. doi: 10.1093/icb/icm039. Epub 2007 Jun 6. PMID: 21672871.

Toyota K, Hiruta C, Ogino Y, Miyagawa S, Okamura T, Onishi Y, Tatarazako N, Iguchi T. Comparative Developmental Staging of Female and Male Water Fleas Daphnia pulex and Daphnia magna During Embryogenesis. Zoolog Sci. 2016 Feb;33(1):31–7. doi: 10.2108/zs150116. PMID: 26853866.

Westphal GH, Stewart Merrill TE. Partitioning variance in immune traits in a zooplankton host-Fungal parasite system. Ecol Evol. 2022 Dec 19;12(12):e9640. doi: 10.1002/ece3.9640. PMID: 36545366; PMCID: PMC9763022.

Zhao H, Chen Z, Li H, Zhao YH, Wang Q, Li WW. Suppressed COP9 signalosome 5 promotes hemocyte proliferation through Cyclin E in the early G1 phase to defend against bacterial infection in crab. FASEB J. 2022 May;36(5):e22321. doi: 10.1096/fj.202101710RRRR. PMID: 35429011.

