## Supplementary Tables and Figures for "Circulating hemocytes continue to proliferate throughout lifespan in *Daphnia*"

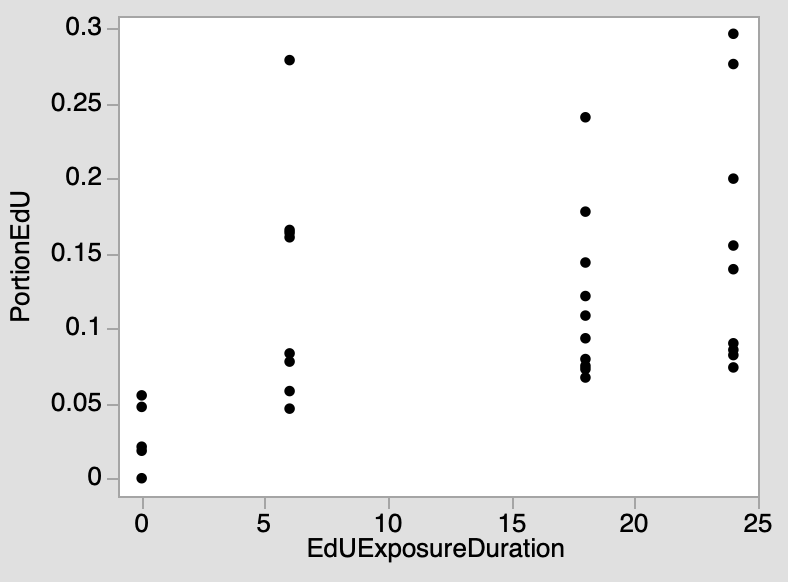

Fig. S1. Portion of hemocytes with EdU staining after 0 – 24 hours of exposure to EdU.

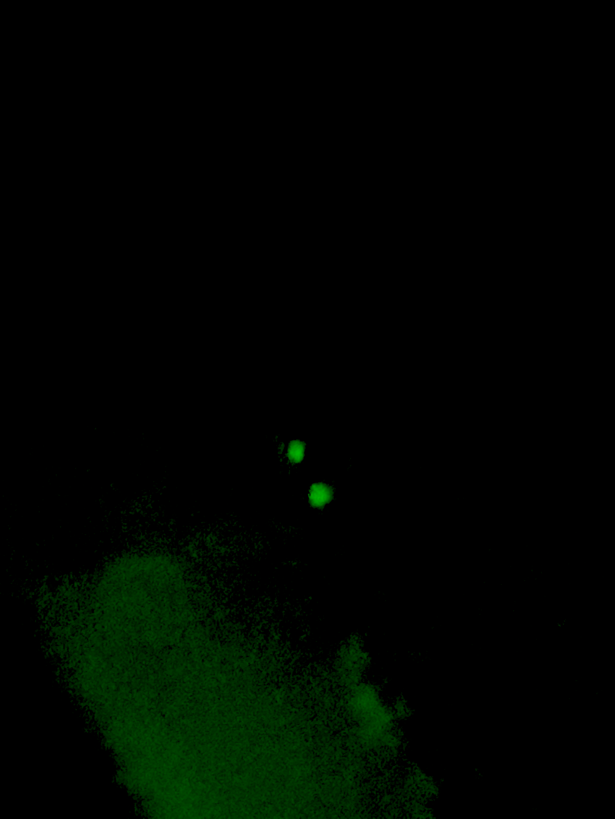
A

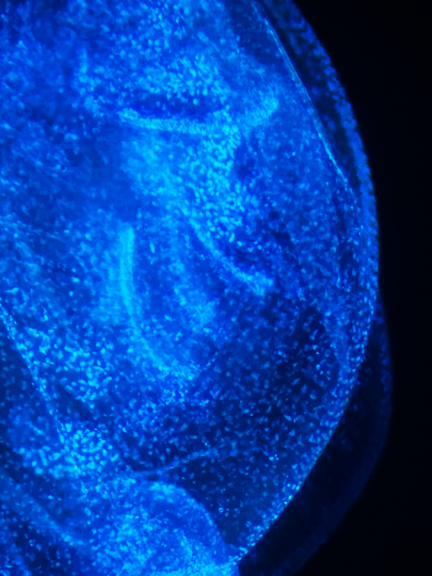

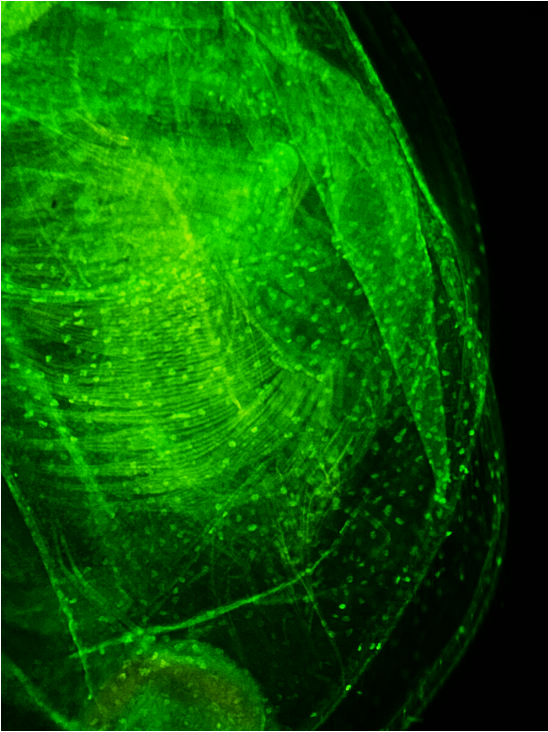
B	C	D

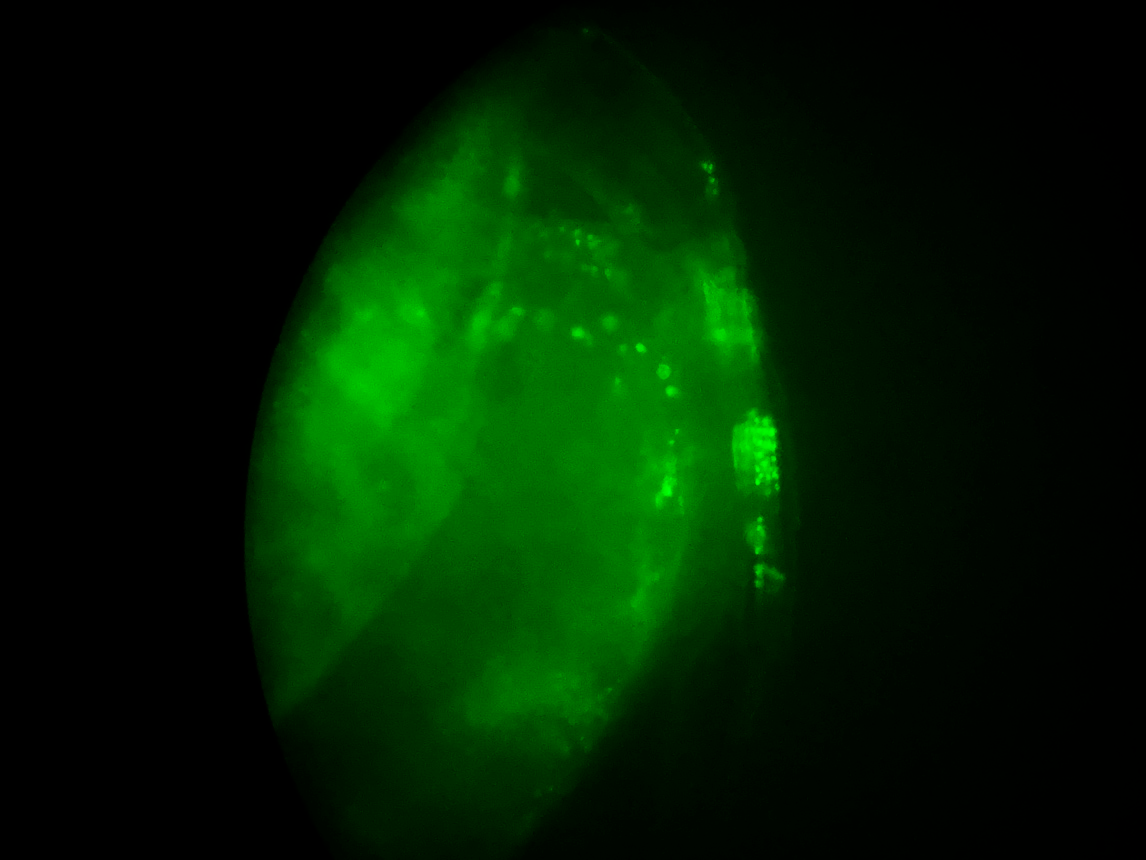

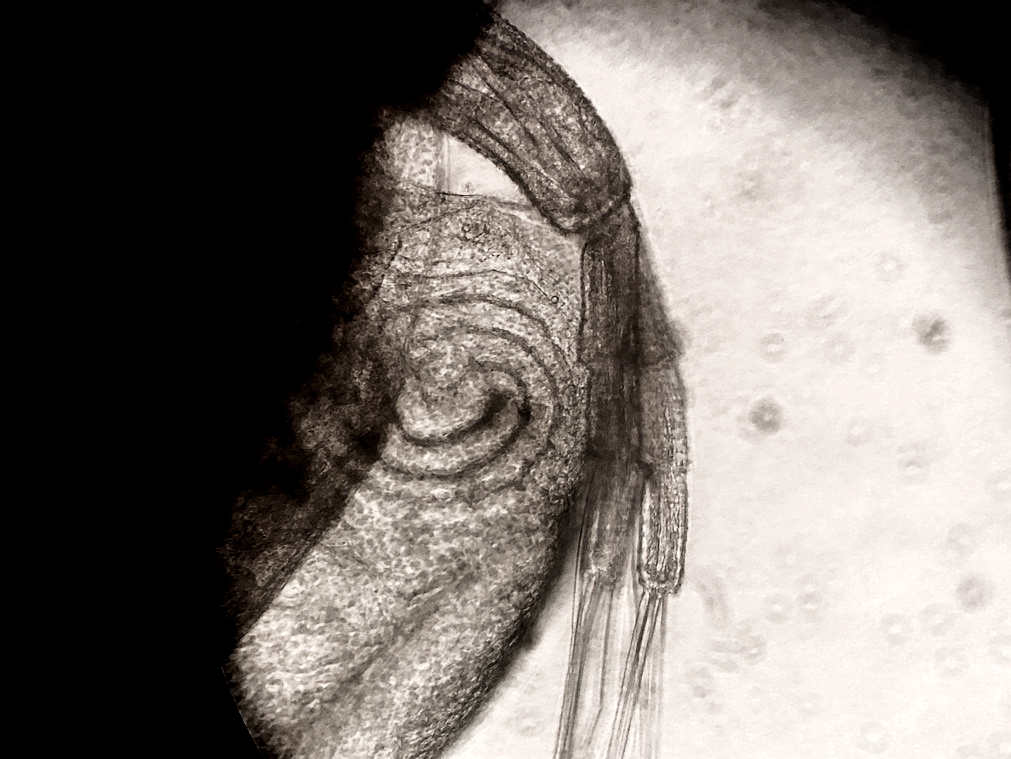

sg

a

e

c

fb

g

Fig. S2. A: a rare example of a pair of EdU-positive nuclei in fat body of an adult *Daphnia* female. B - D: EdU-positive nuclei in identically treated juveniles. B: Antenna and shell gland (EdU fluorescence overlay with bright field); C, D: carapace, fat body and epipodites (C: EdU staining; D: Hoechst). Lowercase letters on B and C: a = antenna-2; sg = shell gland; e = epiodite; fb = fat body; g = gut.

Table S1. Results of REML ANOVA of the effects of age, ovary/molting cycle phase (hours since egg-laying, an ordinal variable), and their interaction on the portion of hemocytes showing EdU staining; longitudinal (top) and common garden (bottom) datasets analyzed separately. Cohorts and replicate sub-cohort nested within cohorts included into the model as random variables. Clutch size could not be included as a main effect in the common-garden analysis due to insufficient data. See main text Table 1 for the same analysis of the complete dataset.

| Longitudinal analysis (2 cohorts, 3 batches) | | | | | | | | |
| --- | --- | --- | --- | --- | --- | --- | --- | --- |
| Source | DF | | DFDen | | F Ratio | | Prob > F | |
| age | 1 | | 10.4 | | 0.340 | | 0.57 | |
| cycle stage | 5 | | 28 | | 1.188 | | 0.34 | |
| age*cycle stage | 5 | | 36 | | 1.395 | | 0.25 | |
| clutch size | 1 | | 36 | | 0.189 | | 0.67 | |
| age*clutch size | 1 | | 35.6 | | 0.376 | | 0.54 | |
| cycle stage*clutch size | 5 | | 31.5 | | 1.120 | | 0.33 | |
| age*cycle stage*clutch size | 5 | | 35.1 | | 1.311 | | 0.28 | |
| Common-garden analysis (5 cohorts, batch 3) | | | | | | | | |
| Source | | DF | | DFDen | | F Ratio | | Prob > F |
| age | | 1 | | 22.5 | | 0.684 | | 0.42 |
| cycle stage | | 5 | | 24.4 | | 0.492 | | 0.78 |
| age*cycle stage | | 5 | | 26.1 | | 0.367 | | 0.87 |
